# FACSanadu: Graphical user interface for rapid visualization and quantification of flow cytometry data

**DOI:** 10.1101/201897

**Authors:** T.R. Bürglin, J. Henriksson

**Affiliations:** Department of Biomedicine, University of Basel, Mattenstrasse 28, 4058 Basel, Switzerland; Wellcome Trust Sanger Institute, Wellcome Trust Genome Campus, Hinxton, CB10 1SA Cambridge, UK

## Abstract

**Motivation:** Flow cytometry is a fundamental technique in cell biology, yet few open source packages are available to analyse these data. Here we describe FACSanadu, an interactive package for rapid visualization and measurement of flow cytometry data. It is the first open source package that can read length profile data from
the COPAS Biosorter.

**Availability and Implementation:** FACSanadu is implemented in Java and uses the Qt framework for display. Binary distributions are made for all major operating systems (Windows, Macintosh, Linux). The source code and documentation is available as free software at http://www.facsanadu.org.

## 1. INTRODUCTION

Flow cytometry is a routine method for analyzing populations of cells. The ease of flow cytometry measurements, in conjunction with antibody markers that allow identification of many cell types, makes it one of the most important tools in biology and for clinical pathology (ref some review). This resulted in a strong need for standardization of file formats (Spidlen et al., 2010). Most instruments from manufacturers come with their own proprietary software, which generally does not allow customizations and makes analysis difficult when data from different instruments are collated. Hence, third party software has become available for data analysis, the two most prevalent software packages being “FlowJo” (http://www.flowjo.com) and “FCS Express” (http://www.denovosoftware.com). But the unavailability of the source code prevents us from adapting it for our purposes. Free analysis packages exist; one is “Flowing” from the Turku Centre of Biotechnology (http://www.flowingsoftware.com). but since it is written in a proprietary programming language (Visual Basic) it is not easy to modify, nor to run on non-windows platforms. A simpler interactive alternative is “FlowPy” (flowpy.wikidot.com), but it also only runs on Windows. For users experienced with the R programming language, the “flowCore” Bioconductor package can handle FCS-files (Hahne et al., 2009). This allows one to use all the powerful statistics of the language, but it is not suitable for the casual flow cytometry users and there is no option to interactively browse the data and define gates. A similar R package able to read the COPAS specific file format exists as well (Shimko & Andersen, 2014) but it only covers the Reflux portion of the instrument. To our knowledge, none of these packages allow data analysis of COPAS Biosorter length profile data. This specialized flow sorter is becoming more popular as larger samples (e.g. *C*. *elegans*, embryos, organoids) are being analyzed more routinely. We hence decided to write our own portable package for flow cytometry analysis, with strong emphasis on fast exploration and ease of use, in particular for COPAS data.

## 2. OVERVIEW OF FACSANADU

FACSanadu is a portable open-source software package implemented in Java using the Qt cross platform application framework (http://www.qt-project.org) and Qt Jambi (http://www.qtjambi.org). This has made it easy to maintain ports for all operating systems, each with the feeling of a modern native user interface. The software can load most common FCS files as well as native files from the COPAS Biosorter.

As expected, FACSanadu can generate scatter plots and histograms of the data. However, to speed up the data processing, we have decided against letting the user manually place out gates on a “worksheet” as commonly found in other programs. Instead, the graphs, upon selection of data, are placed automatically in a compact grid layout; different views are on one axis and datasets on the other axis (Fig 1a). Axis can be changed by simply clicking on the axis label. Only the views and datasets selected in the side bar are displayed. When gates are drawn they are displayed and updated on all views simultaneously. This makes it easy to, for example, show 2 positive and 2 negative controls, and adjust the gating region on all of them simultaneously. The currently displayed views can be exported as PNG files; either as one big image, or split by dataset/view, or as all individual files.

**Fig. 1.**
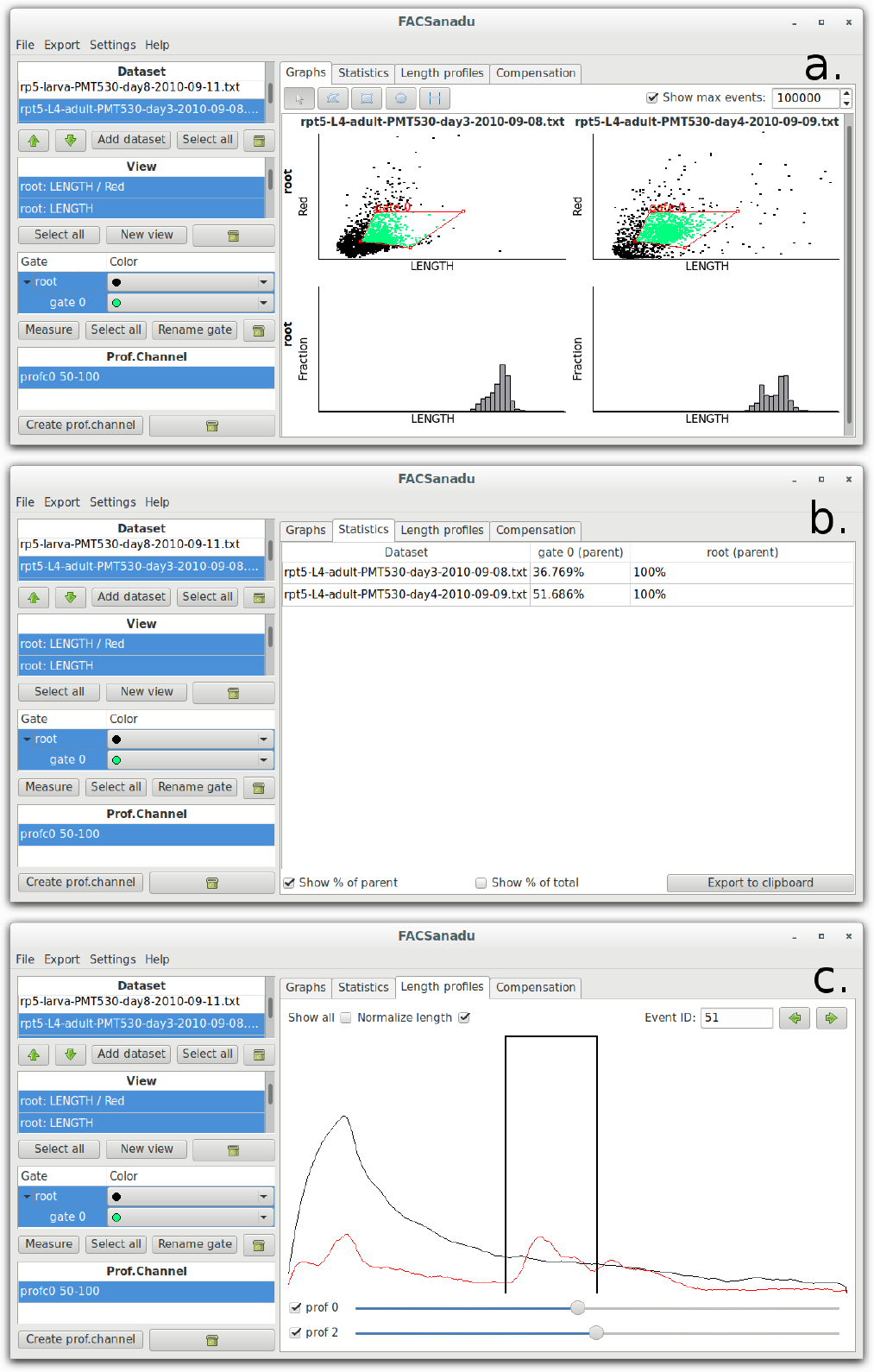
**(A)** Visualization of all views. A grid of plots is made from the current selection of views and datasets. All plots are updated in parallel as gates as drawn or edited. For multiple datasets the gating is done in the background on multiple CPU cores **(B)** Statistical summary table, of the selected datasets and gates, as well as additional intensity measurements **(C)** Display of length patterns of *C. elegans* nematodes as captured on a COPAS Biosorter. A region has been selected from which the integral is provided as a new virtual channel.

After selecting gate regions all the gating statistics can be found on the statistics pane (Fig 1b). A table is generated from the selected datasets and selected gates, with options for showing the fraction of cells from the parent gate, and the total population. Further measurements can be added to each gated population, such as the mean intensity of certain channels. Once all measurements have been set, the table can be copied to the clipboard or exported to CSV.

For data from the COPAS Biosorter, we have added a third pane, which allows the analysis and visualization of length profiles (Fig 1c). The detector of the COPAS Biosorter allows recording of profiles of long objects, for example, the green fluorescent gene expression pattern along the length of the body of *C*. *elegans* larvae. This has previously been used to record chronograms, i.e., the time-evolution of expression patterns of genes along the larvae, and the same automatic alignment algorithm is automatically applied (Dupuy et al., 2007). The length profile pane allows one to manually browse individual objects from the chosen gate, or it can display an ensemble of them. The length profiles can be shown as-is or normalized by length, each channel with an individual scale bar. For interesting regions the user can set a window in which the intensity is measured. This will then appear as a new channel which can be used for gating and statistics as usual.

To allow users to interactively work with large datasets, as well as make full use of modern multi-core CPUs, FACSanadu makes extensive use of threading and background computation. Multiple datasets are always gated in parallel.

## 3. CONCLUSIONS

FACSanadu fills a gap in the flow cytometry analysis toolbox by providing a quick and simple tool for working with flow cytometry data. While we have made an effort to provide the essential functions the user expects from an interactive flow cytometry package, the availability of the source code makes it possible to extend the functions for the needs of particular laboratories. Together with Bioconductor, there is now a complete suite of open source tools for FACS analysis.

## ACKNOWLEDGEMENTS

The authors thank Natalia Kunowska, Valentina Proserpio and Kedar Natarajan for early beta testing of the software as well as suggestions for new features.

